# GABA_B_R agonist baclofen promotes central nervous system remyelination

**DOI:** 10.1101/2022.01.28.478233

**Authors:** Mari Paz Serrano-Regal, Laura Bayón-Cordero, Juan Carlos Chara, Vanja Tepavcevic, Blanca I. Ochoa-Bueno, Carlos Matute, María Victoria Sánchez-Gómez

**Affiliations:** Achucarro Basque Center for Neuroscience, Leioa, Spain; Department of Neurosciences, University of the Basque Country (UPV/EHU), Leioa, Spain; Centro de Investigación en Red de Enfermedades Neurodegenerativas (CIBERNED), Leioa, Spain; Grupo de Neuroinmuno-Reparación, Hospital Nacional de Parapléjicos, SESCAM, Toledo, Spain

**Keywords:** baclofen, oligodendrocyte, GABA_B_ receptor, differentiation, remyelination

## Abstract

Promoting remyelination - the endogenous response by which lost myelin sheaths are regenerated - is considered as a potential neuroprotective strategy to prevent/limit the development of permanent neurological disability in patients with multiple sclerosis (MS). To this end, a number of clinical trials are investigating the potential of existing drugs to enhance oligodendrocyte progenitor cell (OPC) differentiation, the process that fails in chronic MS lesions. As we previously reported that oligodendroglia lineage cells express GABA_B_ receptors (GABA_B_Rs) both *in vitro* and *in vivo*, and that GABA_B_R-mediated signaling enhances OPC differentiation and myelination in *vitro*, here we focused on the remyelinating potential of the best-known GABA_B_R agonist baclofen (Bac), already approved to treat spasticity in MS. We demonstrated that Bac increases myelin protein production following lysolecithin (LPC)-induced demyelination in cerebellar *ex vivo* slices. In addition, Bac administration enhanced OPC differentiation and remyelination in LPC-induced spinal cord lesions in adult mice. Thus, our results suggest that Bac should be considered as a potential therapeutic agent, not only to treat spasticity, but also to improve remyelination in patients with MS.

## Introduction

Multiple sclerosis (MS) is a chronic inflammatory disease of the central nervous system (CNS) characterized by disseminated demyelination (Dendrou et al., 2015). As a consequence of inflammatory demyelination, action potential conduction is disrupted and axons are deprived from metabolic and trophic support, which leads to axonal loss (Lee et al., 2012; Saab et al., 2016), the main correlate of permanent disability in patients with MS (Trapp, 1999). The majority of currently available treatments for MS target CNS inflammation and associated relapses, but do not prevent the development of long-term disability (Kremer et al., 2019a). Thus, the development of therapies to prevent axonal and neuronal loss remains an unmet therapeutic need for patients with MS (Lubetzki et al., 2020). Remyelination is the spontaneous regeneration of myelin that prevents axonal degeneration both in animal models (Irvine and Blakemore; 2008; Mei et al; 2016) and patients with MS (Kornek et al., 2000). However, in most patients, the efficiency of this process decreases significantly with age and disease progression (Franklin and ffrench-Constant, 2017). Therefore, the development of novel treatments that enhance remyelination is a major goal of current MS research, and includes the repurposing of existing drugs (Kremer et al., 2019b).

A block in OPC differentiation (Kotter et al., 2006; Kuhlmann et al., 2008) and lack of myelin sheath formation by surviving mature oligodendrocytes (OLs) have been pointed out as important contributors to remyelination failure in MS (Duncan et al., 2018; Yeung et al., 2019; Heb et al., 2020; Franklin et al., 2021). Therefore, promoting OPC differentiation and improving OL myelination capacity are potential strategies for enhancing myelin repair and preventing neurodegeneration in MS.

Neurotransmitters are important mediators of OPC-neuron communication with a clear influence on OPC behavior (Domercq et al., 2010; Li et al., 2013; Fannon et al., 2015; Zonouzi et al., 2015; Hamilton et al., 2017; Serrano-Regal et al., 2020a). As OPCs receive both excitatory and inhibitory synaptic inputs, mediated by glutamate and γ-aminobutyric acid (GABA) (Bergles et al., 2000; Lin and Bergles, 2004; Káradóttir et al., 2008; Kukley et al., 2008), these molecules have been identified as key regulators of oligodendroglial maturation and myelination (Fannon et al., 2015; Gautier et al., 2015; Serrano-Regal et al., 2020b; Bai et al., 2021).

Regarding myelin repair, GABAergic signaling through GABA_A_Rs has been associated with remyelination after focal demyelination in the rat *corpus callosum* (Kalakh and Mouihate, 2019), as well as in the caudal cerebellar peduncle (Cisneros-Mejorado et al., 2020). GABA_B_Rs have also been suggested as important modulators of myelination given that GABA_B_R antagonism increased OPC proliferation while decreasing their maturation and the production of myelin-related proteins in the developing rat cingulum (Pudasaini et al., 2021). However, the role of oligodendroglial GABA_B_Rs in myelin regeneration remains to be investigated.

Baclofen (Bac), the best known GABA_B_R agonist, is currently used as a therapeutic agent for spasticity in MS, and can be administered either intrathecally or orally because it crosses the blood-brain barrier (Ertzgaard et al., 2017). We previously reported that GABA_B_R activation by Bac promotes differentiation and myelin-protein expression in rat cortical OPC cultures and myelination in OPC-dorsal root ganglion (DRG) neuron cocultures (Serrano-Regal et al., 2020a). Here, we investigated whether Bac modulates remyelination in demyelinated organotypic cerebellar slices as well as in lysolecithin (LPC)-induced spinal cord lesions in adult mice. Our results demonstrate that Bac stimulates myelin protein production *ex vivo* and enhances remyelination *in vivo*, which suggests that this drug may also be a useful therapeutic agent to stimulate remyelination.

## Materials and methods

### Animals

All experiments were conducted with the approval of the ethical committee of the University of the Basque Country (UPV/EHU). Animals were handled in accordance with the European Communities Council Directive and were housed under standard conditions with a 12h light-dark cycle and *ad libitum* access to food and water. All possible efforts were made to minimize animal suffering and the number of animals used. Sprague Dawley rats, C57BL/6 mice, and transgenic mice expressing fluorescence reporter DsRed under the control of the glial-specific proteolipid protein promoter (PLP-DsRed; Hirrlinger et al., 2005), generously provided by Dr. F. Kirchhoff (University of Saarland, Homburg, Germany), were used in this study.

### Cerebellar organotypic slice culture

Slice cultures were made from cerebella of P5-P7 or P11-day-old Sprague Dawley rats and P11-day-old transgenic PLP-DsRed mice according to previously described procedures (Dusart et al., 1997; Doussau et al., 2017; Tan et al., 2018). Briefly, cerebella were cut with a tissue chopper (Mcllwain) into 350 μm parasagittal slices. Meninges were removed and slices were plated onto 0.4 μm pore size Millicell CM culture inserts (Millipore), containing 2-3 slices each. Rat slices were maintained in six-well plates for 13-15 days and mice cerebellar slices for 11 days in culture medium consisting of 50% basal medium with Earle’s salt (BME), 25% Hank’s Balanced Salt Solution (HBSS), 25% inactivated horse serum (all from ThermoFisher Scientific), 5 mg/ml glucose (Panreac), 0.0025 mM _L_-glutamine (Sigma-Aldrich) and antibiotic-antimycotic solution (100 U/ml of penicillin, 100 μg/ml of streptomycin and 0.25 μg/ml of amphotericin B; ThermoFisher Scientific) at 37°C in a humidified atmosphere with 5% CO_2_. Culture medium was replaced every 2-3 days. Slices were treated with GABAergic drugs starting on the day 2 *in vitro* (Supplementary Table 1). Lysolecithin (LPC)-induced demyelination experiments were carried out in cerebellar slices from P11 animals at day 7 *in vitro* by incubation for 16 h with 0.5 mg/ml LPC (Sigma-Aldrich) (Birgbauer et al., 2004). Treatments were performed at the same time as the LPC-stimulus. Slices were fixed in culture inserts with 4% paraformaldehyde (PFA) solution in phosphate-buffered saline (PBS; pH 7.4) for immunohistochemistry (Supplementary Table 2) or processed for western blot analysis at 4 and 6 days after treatment.

### Optic nerve-derived organotypic slice culture

Cultures were obtained from optic nerves of P11-day-old transgenic PLP-DsRed mice. Optic nerves together with the retina were extracted in order to maintain tissue organization and cellular connections. Meninges and residual tissue were removed in supplemented (2 μl/ml gentamicin, 1 mg/ml bovine serum albumin, BSA and 2 mM _L_-glutamine) HBSS under the microscope, and the optic nerve-retina units were maintained in 0.4 μm pore size Millicell CM culture inserts (Millipore), containing one unit each. Explants were maintained in six-well plates for 3 days in the culture medium as described above for cerebellar organotypic cultures and in the same conditions. To favor appropriate feeding of the optic nerve-retina unit, 50 μl of culture medium were added directly over the tissue (Azim and Butt, 2011). GABA_B_R specific agonist baclofen (100 μM) was added to the medium immediately after plating and maintained for 3 days with daily renewal. Optic nerves without retina were fixed with 4% PFA in PBS and whole-mounted on slides with Prolong™ Gold antifade (Invitrogen).

### EdU labeling and detection

5-ethynyl-2’-deoxyuridine (EdU; Invitrogen) (10 μM) was added to the organotypic medium at day 5 *in vitro* and left for 48 h, to label proliferating cells. EdU was revealed using Click-iT Alexa Fluor 647 Imaging Kit according to the manufacturer’s instructions (Invitrogen).

### Demyelinating lesion induction

Demyelinating lesions were induced in the spinal cord of 10-week-old female C57BL/6 mice by a stereotaxic injection of 0.5 μl of 1% LPC (Sigma-Aldrich) in sterile 0.9% NaCl solution, as previously described (Tepavcevic et al., 2014). Mice were anesthetized by intraperitoneal injection (i.p.) of a solution of ketamine (90 mg/kg; Fatro)/xylazine (20 mg/kg; Calier). Buprenorphine (0.1 mg/kg; Dechra) was subcutaneously administered as postoperative analgesic treatment. Daily i.p. injections of vehicle (saline solution) or baclofen (8 mg/kg) were performed from 5-12 days post lesion (dpl). Mice were sacrificed at 12 dpl or 16 dpl, and the tissue was processed for immunohistochemical (IHC) or transmission electron microscopy (TEM) analysis, respectively.

### Perfusion and tissue processing

Mice were euthanized with ketamine/xylazine and transcardially perfused with 2% PFA solution in PBS for IHC analysis or 4% glutaraldehyde in 0.1 M PB for TEM studies. For IHC analysis, spinal cords were postfixed with the same PFA solution, cryoprotected in 15% sucrose solution (Panreac) and frozen in 7% gelatin (Sigma-Aldrich)/15% sucrose solution in PBS. Samples were cut using a cryostat CM3050 S (Leica) to obtain 12 μm-thick coronal sections. For TEM studies, spinal cords were postfixed overnight, washed in 0.1 M PB, and cut into 2 mm-thick blocks. The tissue was postfixed in 1% osmium solution in 0.1 M PB, dehydrated and embedded in epoxy resin (Sigma Aldrich). Semithin (1 μm-thick) and ultrathin (55 nm-thick) sections were cut with an ultramicrotome RMC Boeckeler.

### Immunochemistry

Cerebellar slices were washed in PBS, permeabilized and blocked in 4% goat serum and 0.1% Triton X-100 in PBS (blocking buffer) for 1 h and incubated overnight at 4ºC with primary antibodies (Supplementary Table 2). Slices were washed in PBS with 0.1% Triton X-100 and incubated with Alexa fluorophore-conjugated secondary antibodies (1:400; Invitrogen) in blocking buffer for 1h at RT. Slides with cryostat sections were air-dried for 1 h, rehydrated in Tris buffer saline (TBS; 20 mM Tris and 1.4 M NaCl in dH_2_O; pH 7.6) and pre-treated with absolute ethanol (Sharlab) for 15 min at -20ºC. For APC and Olig2 immunostaining, antigen retrieval was performed by heating the sections in low-pH retrieval buffer (Vector Laboratories) for 45 sec using a microwave. After washing, samples were incubated in blocking buffer solution (1% BSA, 5% goat serum and 0.1% Triton X-100) for 30 min at RT, and then with the primary antibodies diluted in blocking buffer overnight at 4ºC (Supplementary Table 2). Sections were washed in TBS, and incubated with Alexa fluorophore-conjugated secondary antibodies (1:500; Invitrogen) in blocking solution for 1h at RT. Cell nuclei were counterstained with DAPI (4 μg/ml, Sigma-Aldrich) and sections were mounted with Fluoromount-G (SouthernBiotech).

### Image acquisition and analysis

Images from cerebellar organotypic slices and optic nerve explants were acquired using Zeiss LSM800 and/or Leica TCS SP8 laser scanning confocal microscopes. Cells in cerebellar slices were counted blindly along the z-stack using a 20x objective in Leica TCS SP8 confocal microscope. At least 3 different fields from 2 slices per experiment were analyzed by using LAS AF Lite software (Leica). The fluorescence signal corresponding to the PLP-DsRed OLs was quantified by ImageJ software and data were expressed as arbitrary units of fluorescence for each experimental situation. Images from spinal cord sections were collected using 40x and 63x objectives in Leica TCS SP8 confocal microscope and imported to *ImageJ* software. Area lacking myelin basic protein (MBP) staining within the *dorsal funiculus* of the spinal cord (area of demyelination) was delimited as region of interest (ROI) and measured. Cells positive for the markers of interest were counted from at least 3 different slices per animal. Results are presented as percentage or total number of positive cells per lesion area measured. Same settings were kept for all samples (control and treated) belonging to a specific experiment. All images are shown as projections from z-stacks. For TEM studies, semithin sections were used to identify the lesion area after staining with Richardson’s blue, and ultrathin sections were contrasted by incubation in a 4% uranyl acetate and lead citrate solution for its visualization in a Philips CM200 transmission electron microscope. Remyelination was determined as the percentage of OL and SC-remyelinated axons within the total numbers of axons initially demyelinated (those remyelinated+those demyelinated).

### Western blot

After treatments, cerebellar slices were directly resuspended in sodium dodecyl sulfate sample buffer on ice to enhance the lysis process and avoid protein degradation. Samples were boiled at 99ºC for 8 min, size-separated by sodium dodecyl sulfate polyacrilamide gel electrophoresis (SDS-PAGE) in 4-20% Criterion TGX Precast gels and transferred to Trans-Blot Turbo Midi PVDF Transfer Packs (Bio-Rad, Hercules, USA). Membranes were blocked in 5% BSA (Sigma-Aldrich) in Tris-buffered saline/ 0.05% Tween-20 (TBS-T) and proteins were detected with specific primary antibodies (Supplementary Table 3). Membranes were incubated with horseradish peroxidase-conjugated secondary antibodies (1:2000; Sigma-Aldrich) and were developed by using an enhanced chemiluminescence detection kit according to the manufacturer’s instructions (Supersignal West Dura or Femto; ThermoFisher Scientific). Protein bands were detected with a ChemiDoc XRS Imaging System (Bio-Rad) and quantified by volume using *ImageLab* software (version 3.0; Bio-Rad).

### Statistical analysis

All data are presented as mean ± SEM. Statistical analyses were performed using *GraphPad Prism* statistical software (version 8.0; GraphPad software). Comparisons between multiple experimental groups were made using one-way analysis of variance (ANOVA) followed by Tukey’s *post hoc* test. For comparisons between two groups, we used the two-tailed Student’s *t*-test assuming equal variance. In all instances, statistical differences were considered significant where p < 0.05. All the images shown represent the data obtained from at least three independent experiments.

## Results

### Baclofen treatment upregulates myelin protein levels in organotypic slice cultures

We first validated the role of the GABAergic signaling in regulating oligodendroglial differentiation and myelination in murine organotypic cultures obtained from P5-P7 rats. We investigated GABA_B1_ and GABA_B2_ receptor-subunit expression during myelination *ex vivo*, and found that oligodendroglial cells - labeled using anti-Olig2 antibody -, and more specifically mature OLs - labeled using anti APC antibody - express the two GABA_B_R subunits (**Fig. 1A, 1B**), as we previously observed in isolated cultured OLs (Serrano-Regal et al., 2020a). We then assessed by western blot whether GABA agonists modulate myelin-protein expression levels in these preparations. Exposure to 100 μM GABA or 100 μM muscimol (Mus) (GABA_A_R specific agonist) did not change the levels of expression of myelin-associated glycoprotein (MAG), 2’,3’-cyclic nucleotide-3’-phosphodiesterase (CNPase) and MBP, compared with control slices (Supplementary Figs. 1 and 2). However, treatment with 100 μM Bac (**Fig. 1C**) induced a significant increase in the expression of MAG and MBP myelin proteins (2.0 ± 0.34 Bac *vs* 1.0 ± 0.19 control for MAG, **Fig. 1D;** and 1.29 ± 0.14 Bac *vs* 1.0 ± 0.12 control for MBP, **Fig. 1F**), together with a non-significant increase in the expression of CNPase (1.23 ± 0.11 Bac *vs* 1.0 ± 0.14 control for CNPase, **Fig. 1E**). To check if this effect of Bac was associated with changes in oligodendroglial proliferation, cerebellar organotypic cultures were exposed to EdU for 48 h, in the absence or presence of GABA or Bac (100 μM). Neither GABA nor Bac modified the percentage of mature OLs (APC^+^Olig2^+^) among total oligodendroglial cells, nor Olig2^+^ cells that underwent proliferation in this time period (Olig2^+^Edu^+^) **(Fig. 1G, H**, and **I)**, suggesting that GABA_B_R activation promotes myelin generation by mature OLs without affecting OPC proliferative capacity. Additionally, we examined the effect of Bac directly over mature OLs in optic nerve explants of transgenic PLP-DsRed mice (**Fig. 1J**). Quantification of PLP-DsRed-fluorescent signal (**Fig. 1K**) revealed a significant increase in those optic nerves treated with Bac compared to controls (27.82 ± 2.27 for Bac *vs* 19.62 ± 2.59 for control; **Fig. 1L**). Together, these results show that Bac enhances myelin protein production in murine organotypic cultures and in optic nerve explants, confirming our previous observations in isolated OLs (Serrano-Regal et al., 2020a) in a complex environment more similar to physiological conditions.

**Figure 1.**
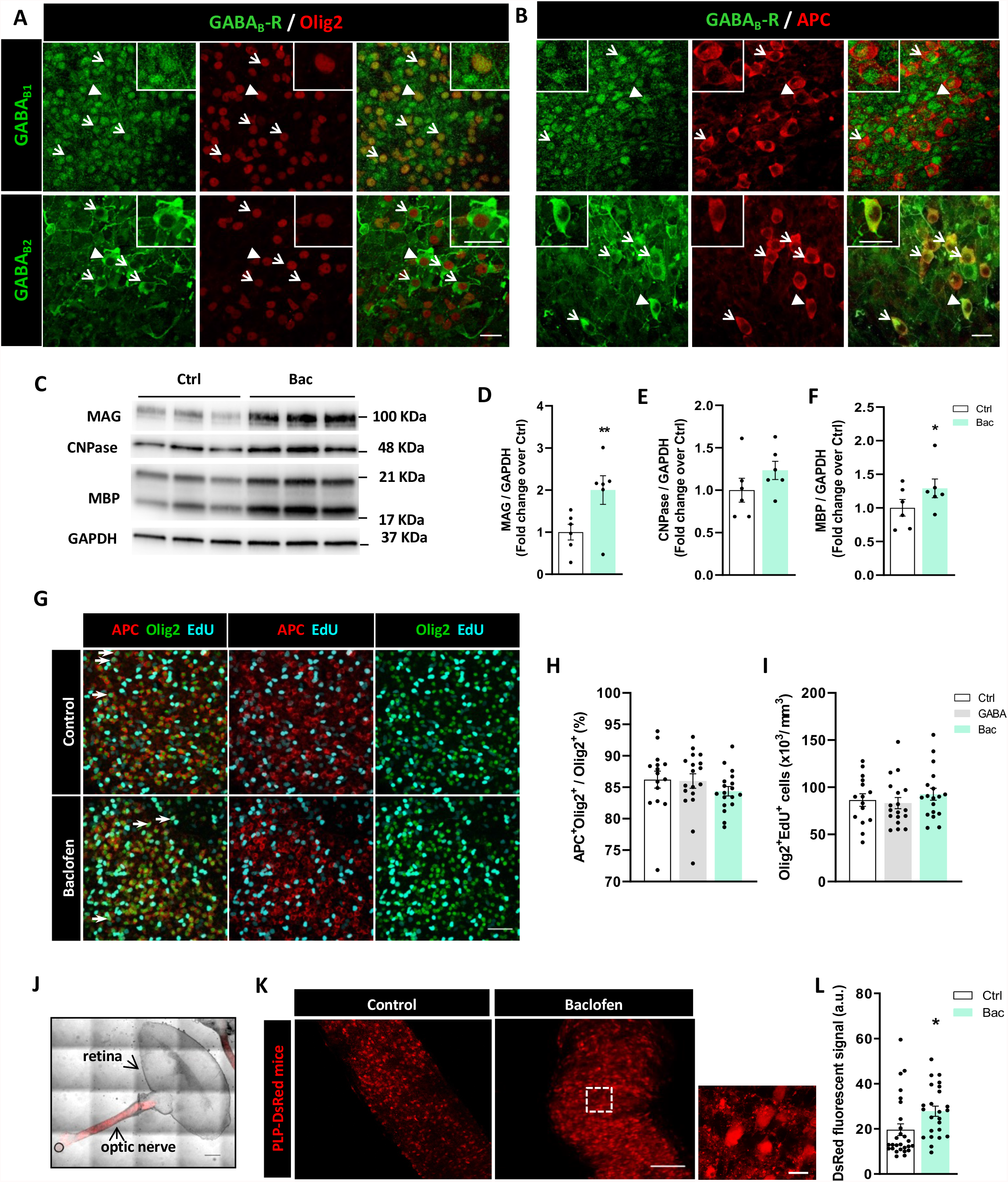
Baclofen increases myelin-related protein synthesis in organotypic cultures without altering the proliferation ratio of oligodendroglial lineage. **A)** Oligodendroglial cells, distinguished as Olig2^+^ cells (red), are positive for GABA_B1_ and GABA_B2_ subunits (green) of GABA_B_Rs in cerebellar slices of P5-P7 rats. **B)** Mature OLs, identified as APC^+^ cells (red), express GABA_B1_ and GABA_B2_ subunits (green) of GABA_B_Rs in the same preparations. Arrows indicate double-stained cells and arrowheads show the cell magnified in the corresponding inset. Scale bars = 20 μm. **C)** Representative western blot image showing expression of MAG, CNPase and MBP proteins in control and baclofen-treated cerebellar slices. Quantification of MAG **(D)**, CNPase **(E)** and MBP **(F)** expression normalized to GAPDH values. *p<0.05 and ** p<0.01 *vs* control; paired Student’s *t*-test. **(G)** Representative images showing immunofluorescence of mature OLs (APC^+^, red) and total oligodendroglial cells (Olig2^+^, green) co-labeled with EdU (cyan) to identify mature OLs (APC^+^) and Olig2^+^ cells in cerebellar slices in the indicated condition. Arrows indicate mature OLs (APC^+^Olig2^+^) or newly generated oligodendroglial cells (Olig2^+^EdU^+^). Scale bar = 50 μm. Quantification of **(H)** percentage of mature OL from total oligodendroglial cell pool (APC^+^Olig2^+^/Olig2^+^) and **(I)** density of newly formed oligodendroglial cells (Olig2^+^EdU^+^), in the indicated conditions. One-way ANOVA followed by Tukey’s post-test. **(J)** Optic nerve-retina unit from P11 PLP-DsRed transgenic mice. Scale bar = 500 μm. **(K)** Optic nerves explants cultured in control conditions (left) or in presence of baclofen (right) showing DsRed fluorescent signal. Scale bars = 100 μm; higher magnification scale bar = 315 μm. **L)** Quantification of DsRed fluorescent signal in control and treated optic nerve explants. *p<0.05 *vs* control; unpaired Student’s *t*-test. At least 3 independent experiments were included in the analysis.

### GABA_B_R activation enhances myelin protein expression during remyelination *ex vivo*

Since Bac promoted myelin protein synthesis in organotypic slices, we next studied the impact of GABA_B_R activation under experimental conditions mimicking damage to myelin. P11 rat-derived cerebellar slices were maintained for 7 days to allow myelination *ex vivo* and then exposed to LPC for 16 h. In a first set of experiments, slices were daily treated with GABA (100 μM) or Bac (100 μM) for 6 days after LPC exposure (Supplementary figure 3A and B) and MAG and MBP proteins were analyzed by western blot. We found that LPC induced a strong decrease of both proteins (0.99 ± 0.11 LPC *vs* 1.44 ± 0.11 control for MAG, and 1.00 ± 0.08 LPC *vs* 1.50 ± 0.09 control for MBP), while Bac treatment post-LPC significantly increased MAG levels (1.44 ± 0.11 Bac *vs* 0.99 ± 0.11 LPC; Supplementary figure 3C) without changing MBP levels (Supplementary figure 3D). In view of that, we then investigated if previous application of these agonists was more effective in fostering remyelination. Thus, we exposed cerebellar slices to LPC in the presence of GABA or Bac (100 μM) and treatments were maintained for 6 days after LPC removal (**Fig. 2A**). MAG and MBP levels were analyzed again by western blot (**Fig. 2B**), and, under this experimental paradigm, we observed a significant increase in the expression levels of both myelin proteins (2.75 ± 0.35 GABA and 2.81 ± 0.26 Bac *vs* 1.0 ± 0.26 LPC for MAG, and 3.39 ± 0.52 GABA and 2.82 ± 0.29 Bac *vs* 1.0 ± 0.37 LPC for MBP; **Fig. 2C** and **D**, respectively).

**Figure 2.**
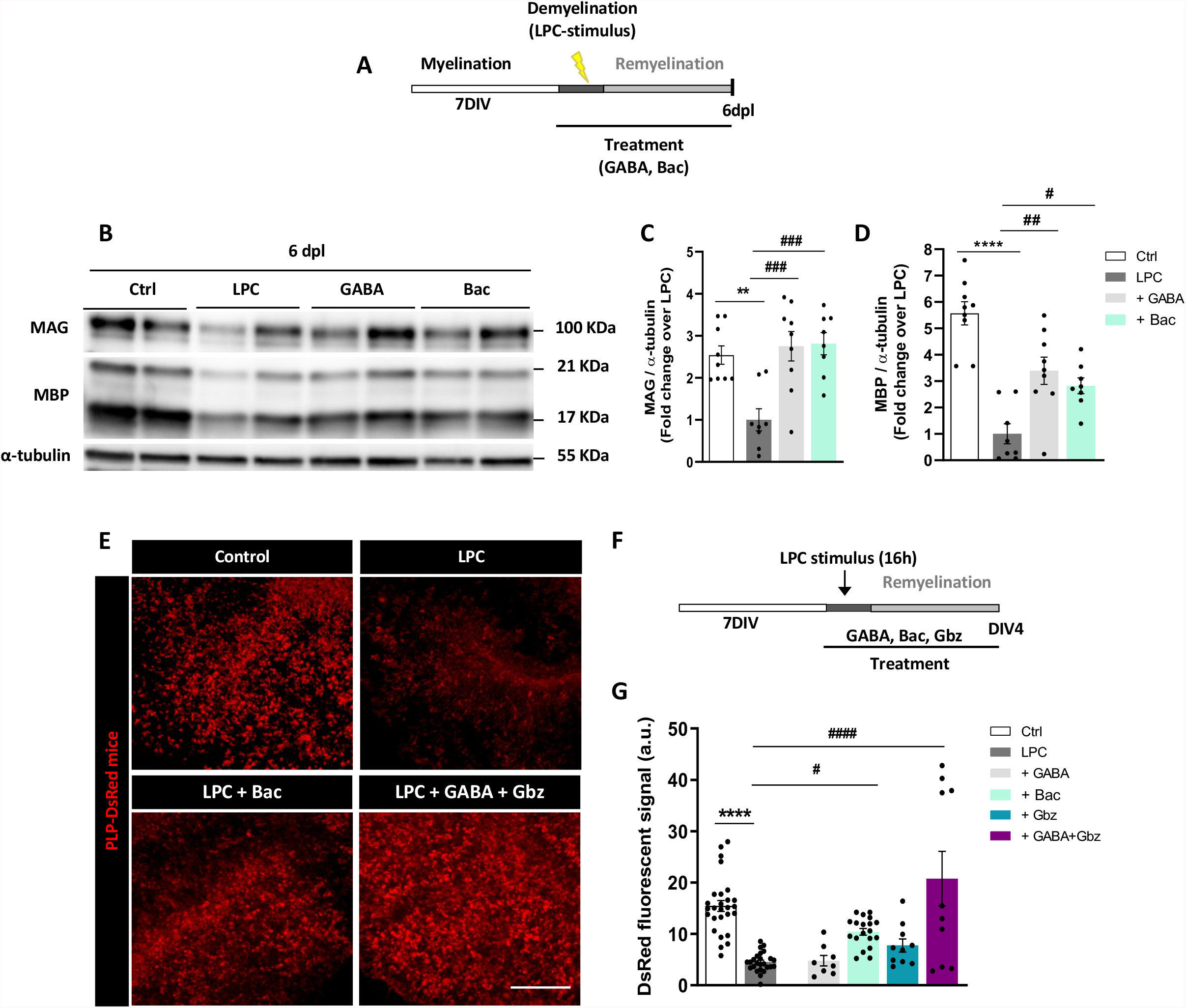
GABA_B_Rs modulate remyelination in LPC-treated cerebellar organotypic slices. **(A)** Time course showing the experimental design in LPC-induced demyelination in cerebellar organotypic cultures obtained from P11 rats. **(B)** Representative western blot image showing in duplicates the effect of GABA and baclofen in modulating myelin-related protein restoration in LPC-treated organotypic cultures following the paradigm shown in A. **(C, D)** Quantification of MAG (C) and MBP (D) levels in indicated conditions. **p<0.01 and ****p<0.0001 *vs* control, ^##^p<0.01 and ^###^p<0.001 *vs* LPC; one-way ANOVA followed by Tukey’s post-test. **(E)** Representative images of cerebellar slices from P11 PLP-DsRed transgenic mice showing DsRed fluorescence in indicated conditions. Scale bar = 100 μm. **(F)** Treatments were added to the slices in conjunction with LPC for 16 h and maintained thereafter for 4 days after. GABA_B_Rs were selectively activated with baclofen or with GABA plus the GABA_A_R antagonist gabazine. **(G)** Quantification of DsRed fluorescent signal in indicated conditions. ****p<0.0001 *vs* control, ^#^p<0.05 and ^####^p<0.001 *vs* LPC; one-way ANOVA followed by Tukey’s post-test.

To confirm whether GABA_B_R-mediated signaling increases OL differentiation and myelin protein production *ex vivo*, we used cerebellar slices from P11 PLP-DsRed transgenic mice, taking advantage of the PLP-associated endogenous fluorescence. We maintained the slices for 7 days and we next applied LPC in combination with drug treatments, added for 4 days. We applied GABA (100 μM), Bac (100 μM) and GABA plus gabazine (50 μM) - a GABA_A_R antagonist -, in order to study the effect of GABA directly over GABA_B_Rs, or gabazine alone (50 μM) (**Fig. 2E, 2F**). As shown in **Fig. 2G**, the PLP-DsRed fluorescent signal increased significantly in Bac- and GABA plus gabazine-treated slices compared to those exposed to LPC without treatment (10.42 ± 0.65 Bac and 20.74 ± 5.32 GABA plus gabazine *vs* 4.5 ± 0.38 LPC; **Fig. 2G**), indicating that GABA_B_ receptor stimulation in these slices increases PLP production. Overall, these results suggest that GABA_B_R activation with Bac favors remyelination *ex vivo* in cerebellar organotypic slices.

### Baclofen administration promotes OPC differentiation in adult mouse CNS

We then assessed the effect of Bac administration on CNS remyelination *in vivo*. Demyelination was induced by LPC injection in the *dorsal funiculus* of young adult mice, and 5 days later daily i.p. injections of vehicle or Bac (8 mg/kg/day) were initiated and administered during 7 days. OPC differentiation was investigated at 12 dpl, given that the peak of OPC differentiation occurs during the second week post demyelination (**Fig. 3A**).

**Figure 3.**
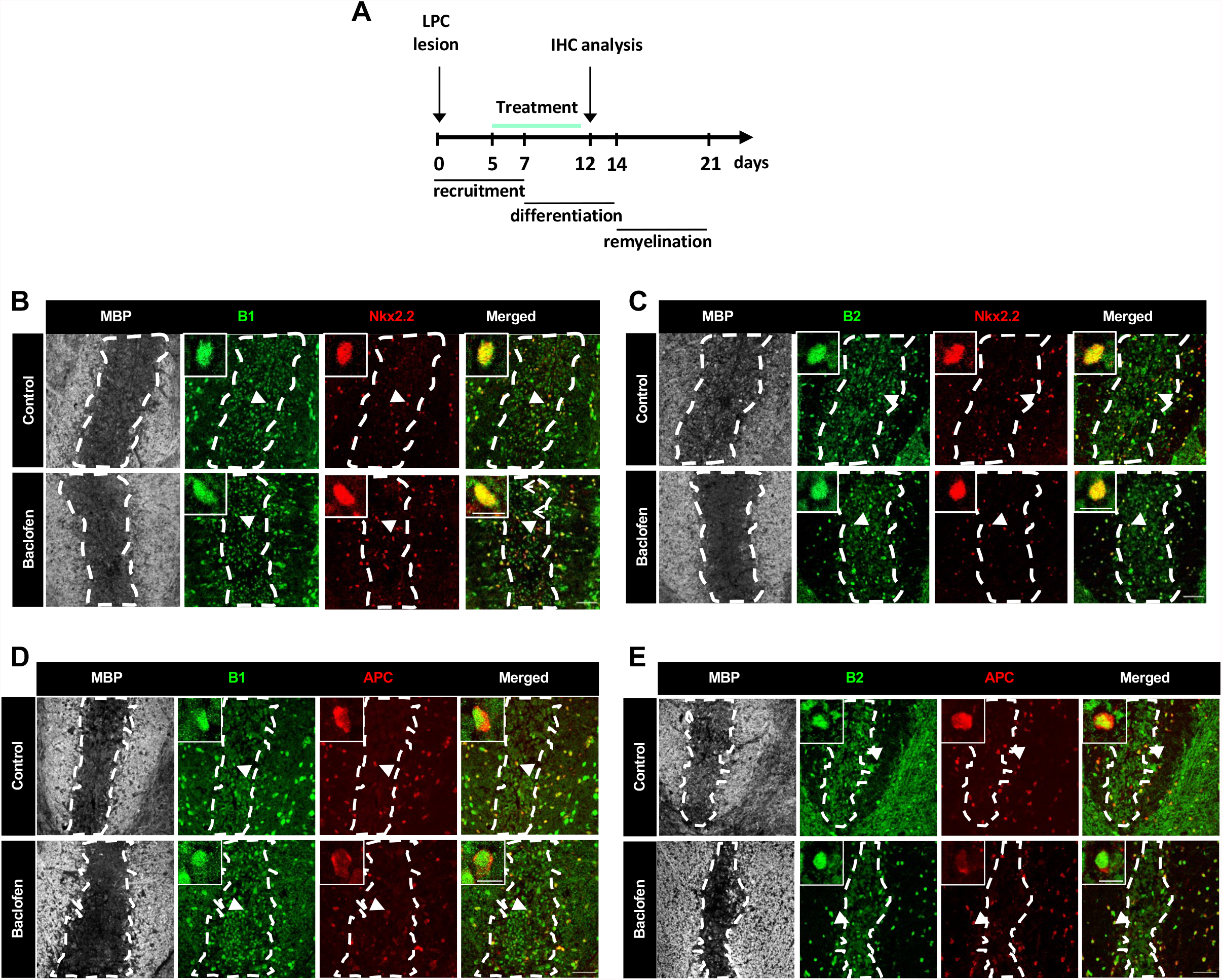
GABA_B_ receptors are expressed by OPCs and mature oligodendrocytes from the spinal cord of LPC-lesioned mice. **(A)** Diagram representing the time course of the studies in LPC-induced demyelinated murine spinal cords. Confocal images showing OPCs (Nkx2.2^+^, red) (**B, C**) and mature OLs (APC^+^, red) (**D, E**) expressing GABA_B1_ (green; B, D) and GABA_B2_ (green; C, E) subunits of GABA_B_Rs in the *dorsal funiculus* of the spinal cord of control (top) and baclofen-treated (bottom) LPC-injected mice. White dash line indicates lesion border. Arrows indicate Nkx2.2^+^/APC^+^ cells expressing GABA_B_Rs subunits. Arrowheads point at cells shown at higher magnification in each photograph. Scale bars = 50 μm. Higher magnification = 10 μm.

We first verified the expression of GABA_B_R subunits in OPCs and mature OLs in normal spinal cord tissue (Supplementary **figure 4A** **and** **B**, respectively) and in demyelinated lesions (**Fig. 3B, C, D** and **E**) using GABA_B_R-subunit specific antibodies, the OPC marker Nkx2.2, and the mature OL marker APC. MBP staining was used to identify the area of demyelination. At 12 dpl, GABA_B1_ and GABA_B2_ subunit expression was observed in both OPCs and mature OLs in the control spinal cord (Suplementary **figure 4A** **and B**, respectively), as well as in recruited OPCs and mature OLs within the lesions of vehicle- (**Fig. 3B, C, D, E**, top) and Bac-treated mice (**Fig. 3B, C, D, E**, bottom).

Then, we explored the changes induced by Bac administration in OPC or microglia/macrophage numbers at 12 dpl using anti-PDGFRα antibody as OPC marker and anti-Iba1 antibody as microglia/macrophage marker (**Fig. 4A**). Quantification of PDGFRα^+^ and Iba1^+^ cells per mm^2^ in control *vs* Bac-treated mice did not reveal any variation (401.2 ± 72.4 Bac *vs* 461.3 ± 49.93 vehicle for PDGFRα^+^ cells; **Fig. 4B** and 5204 ± 1390 Bac *vs* 6180 ± 258.5 vehicle for Iba1^+^ cells; **Fig. 4C**). To corroborate these results, and investigate the effect of Bac treatment over astrogliosis, we used anti-Nkx2.2 antibody for OPC labeling and anti-GFAP antibody as astrocyte marker (**Fig. 4D**). Similar to previous results, we confirmed the lack of significant differences comparing vehicle *vs* Bac-treated animals in the analysis of Nkx2.2^+^ cells per mm^2^ (437.4 ± 46.8 Bac *vs* 496.5 ± 65.5 control) neither in GFAP stained area (0.024 ± 0.004 mm^2^ Bac *vs* 0.033 ± 0.006 mm^2^ control). Finally, we investigated whether Bac administration accelerates OPC differentiation inside the lesion by analyzing the presence of mature OLs (APC^+^Olig2^+^) relative to the total number of oligodendroglial cells (Olig2^+^) (**Fig. 4D**). The percentage of APC^+^ among total Olig2^+^ cells was significantly increased in Bac-treated LPC-injected mice (41.91 ± 2.29 % Bac *vs* 27.95 ± 1.46 % vehicle; **Fig. 4E**), without altering the total numbers of Olig2^+^ cells (928.0 ± 100.7 Bac *vs* 814.8 ± 50.13 vehicle; **Fig. 4F**). Therefore, Bac treatment promotes differentiation of OPCs in LPC-induced demyelinating lesions.

**Figure 4.**
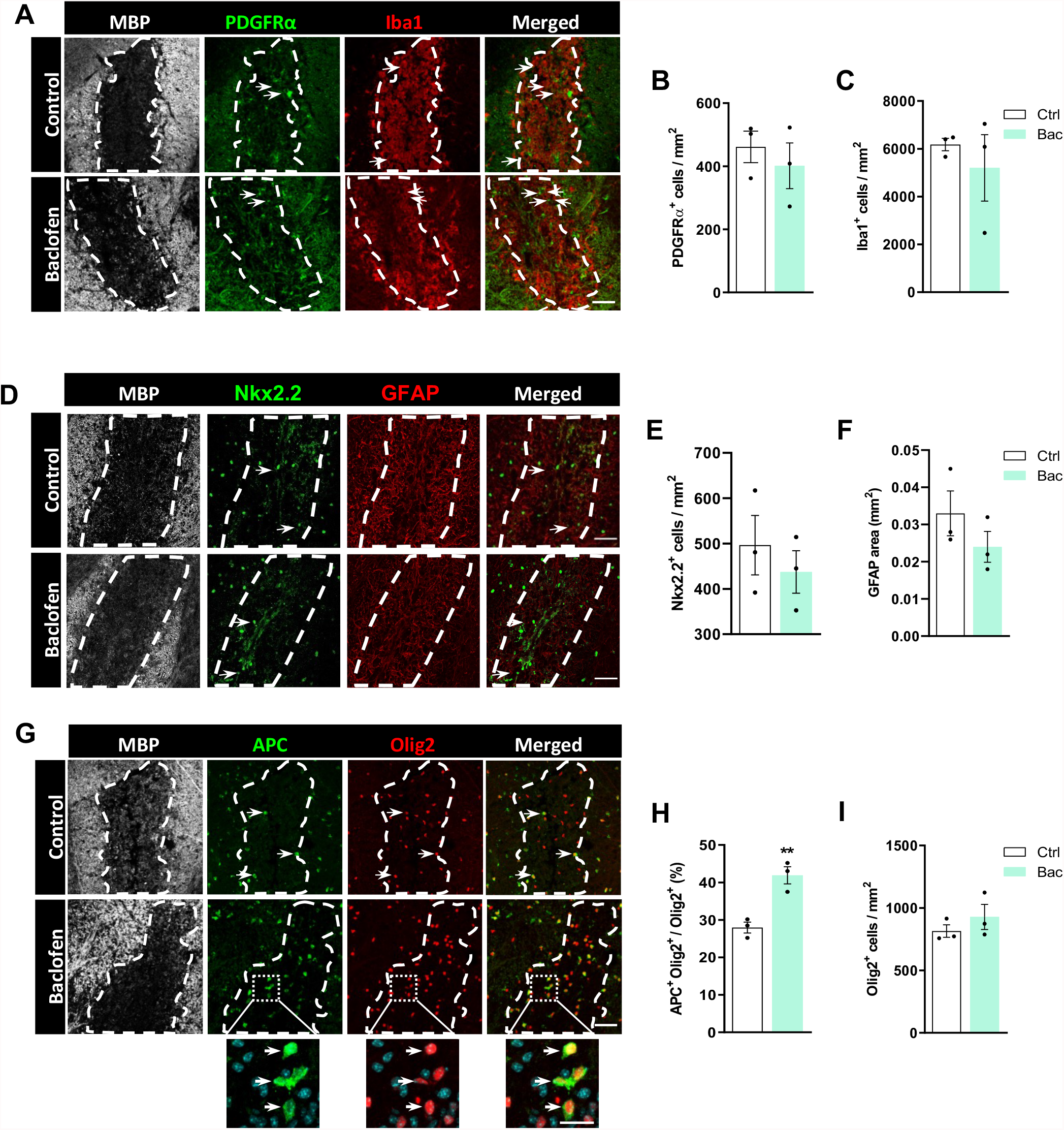
GABA_B_ receptor activation by baclofen promotes oligodendrocyte differentiation but does not impact on OPC nor microglia/macrophague population in LPC-induced demyelinated spinal cords. **(A)** Spinal cord sections of LPC-injected control (top) and baclofen-treated (bottom) mice immunostained with anti-MBP (grey), anti-PDGFRα (green) and anti-Iba1 (red) antibodies. Dapi was used to identify cell nuclei. White dash line indicates lesion border. Quantification of number of PDGFRα^+^ cells **(B)** and number of Iba1^+^ cells per mm^2^ **(C)** in LPC-injected mice. **(D)** Spinal cord sections of LPC-injected control (top) and baclofen-treated (bottom) mice immunostained with anti-MBP (grey), anti-Nkx2.2 (green) and anti-GFAP (red) antibodies. Dapi was used to identify cell nuclei. White dash line indicates lesion border. Quantification of number of Nkx2.2^+^ cells **(E)** and number of GFAP stained area as mm^2^ **(F)** in LPC-injected mice. **(F)** Spinal cord sections of LPC-injected control (top) and baclofen-treated mice immunostained with anti-MBP (grey), anti-APC/CC1 (green) and anti-Olig2 (red) antibodies. Dapi was used to identify cell nuclei. White dash line indicates lesion border. Histograms showing percentage of APC^+^Olig2^+^ cells from total Olig2^+^ cells **(G)**, and number of Olig2^+^ cells per mm^2^ **(H)** in LPC-injected mice. At least 3 lesion areas from 3 different animals were analyzed. **p< 0.01 *vs* control; unpaired Student’s *t*-test. Scale bars: **A, D** = 50 μm. Higher magnification = 20 μm. Arrows show positive staining.

### Baclofen administration accelerates remyelination

We next analyzed whether Bac administration accelerates remyelination. Vehicle and Bac injections were administered following the same time course as in previous experiments, and the proportion of remyelinated axons was analyzed at 16 dpl (**Fig. 5A**), given that the onset of remyelination in the LPC model takes place at 14 dpl and is near completion at 21 dpl. At this time point, OL remyelination - identified as thin myelin sheaths - was found predominantly around lesion borders, while SC remyelination, a well-recognized feature of the LPC lesion in the spinal cord (Jeffery and Blakemore, 1995), was observed in the lesion center. TEM analysis revealed that the percentage of remyelinated axons (**Fig. 5C**) was higher following Bac treatment (36.95 ± 2.18 % Bac *vs* 22.29 ± 1.85 % vehicle). In particular, the percentage of axons remyelinated by OLs was increased after Bac administration (33.45 ± 3.95 % Bac *vs* 19.75 ± 2.63 % vehicle; **Fig. 5D**), whereas SC-driven remyelination was not significantly altered (3.70 ± 2.02 % Bac *vs* 2.54 ± 1.82 % vehicle; **Fig. 5E**). Thus, Bac injection following LPC-induced demyelination accelerates the regeneration of myelin sheaths.

**Figure 5.**
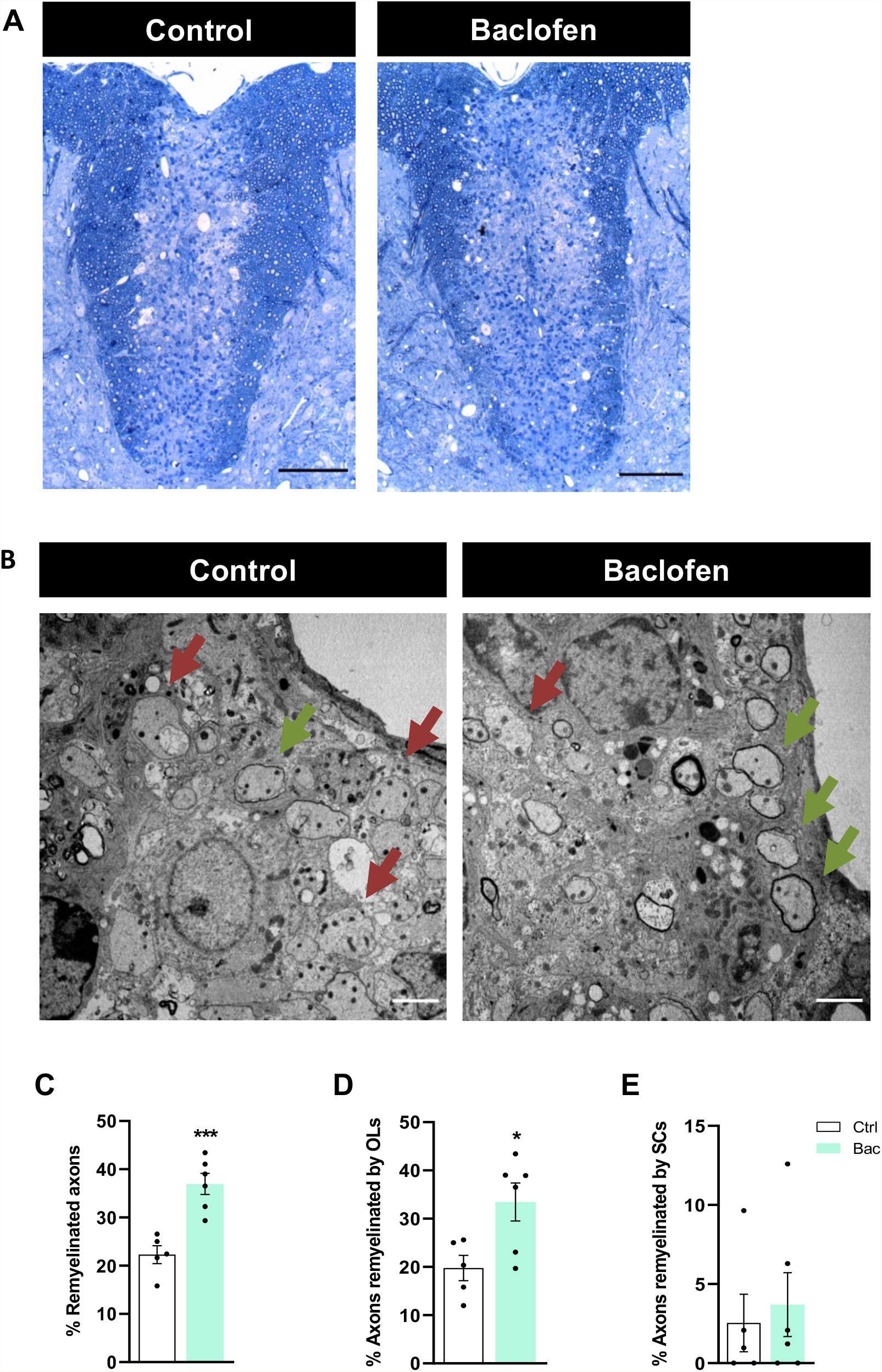
Baclofen treatment promotes remyelination following spinal cord LPC-induced demyelination. **(A)** Representative images of semithin sections of spinal cord control (left) and baclofen-treated (right) mice showing LPC-induced lesions, stained with Richardson’s blue. **(B)** Electron micrographs of ultrathin spinal cord sections showing remyelinated (green arrows) and unmyelinated (red arrows) axons. Histograms showing percentage of total remyelinated axons within the lessions **(C)**, axons remyelinated by OLs **(D)** and axons remyelinated by SCs **(E)**. *p< 0.05 and ***p<0.001 *vs* control; unpaired Student’s *t*-test. Scale bars: **A** = 100 μm; **B** = 2 μm.

## Discussion

Remyelination varies considerably between MS patients, and its presence is associated with higher extent of axonal preservation in the lesions (Kornek et al., 2000) and milder levels of disability (Bodini et al., 2016). As OPC differentiation arrest is an important contributor to failed myelin repair in MS (Kotter et al., 2006; Kuhlmann et al., 2008), promoting OPC differentiation may enhance remyelination and prevent axonal degeneration in the disease, as observed in animal models (Irvine and Blakemore, 2008; Mei et al., 2016). Identifying novel pro-remyelinating agents among already approved drugs is therefore of major interest as it would significantly shorten the time required for bench-to-bedside transition.

We previously showed that GABA_B_R-selective agonist Bac, a drug used for the treatment of spasticity in MS patients (Chisari et al., 2020) stimulates OPC differentiation *in vitro* via GABA_B_R activation (Serrano-Regal et al., 2020a). Interestingly, it has been highlighted that intrathecal Bac administration in MS patients is linked to improved cognitive performance (Farrel et al., 2021). Notably, higher cognitive performance in patients with MS has been associated to quantitative improvements in myelin water imaging (Abel et al., 2020). Here, we show that Bac administration stimulates myelin protein synthesis and remyelination *ex vivo* and *in vivo*.

The addition with GABA and Bac to LPC-treated (demyelinated) control cerebellar slices and optic nerve explants induced an increase in myelin proteins. Moreover, GABA increased PLP levels when applied in conjunction with the GABA_A_R antagonist gabazine.

These results suggest that GABA_B_R activation mediates synthesis of myelin proteins. Importantly, the increase in myelin proteins observed in Bac-treated slices following demyelination strongly suggested that Bac may stimulate remyelination. Therefore, we investigated whether Bac administration can improve remyelination in the LPC-induced model of demyelination *in vivo*.

LPC model of demyelination is characterized by a defined sequence of events: demyelination is completed within 2 days post lesion (dpl), which is then followed by OPC recruitment and proliferation (1^st^ week post LPC injection), OPC differentiation (7-14 dpl), and remyelination (14-21 dpl) (Piaton et al.; 2011; Tepavcevic et al.; 2014; Wegener et al.; 2015). We first demonstrated the expression of GABA_B1_ and GABA_B2_ subunits of GABA_B_Rs by OPCs and newly-generated OLs at 12 dpl, which suggests these cells will be responsive to Bac. To investigate whether Bac treatment influences OPC differentiation, Bac administration was performed starting at 5 dpl, after initiation of OPC recruitment and prior to the onset of differentiation. This treatment resulted in increased percentage of Olig2^+^ cells that express the mature marker APC/CC1 at 12 dpl. Moreover, we did not detect significant changes in OPC numbers (PDGFRa^+^/Nkx2.2^+^ cells), nor in the total oligodendroglial pool (Olig2^+^) at this time point, which suggests that Bac treatment led to an increase in OPC differentiation. Remyelination in LPC lesions starts around 14 dpl. Importantly, Bac-treated mice showed an increase in the percentage of remyelinated axons at 16 dpl, indicative of accelerated remyelination. Thus, Bac-treated mice showed increased OPC differentiation and accelerated remyelination.

Bac administration could increase OPC differentiation (and consequently remyelination) by multiple mechanisms. The first one is via activation of GABA_B_Rs, that, as we showed here, are expressed by oligodendroglial cells in the lesions. This mechanism would be consistent with our previous observations *in vitro* that Bac treatment of OPC cultures increases OPC differentiation, and that this effect is abrogated by application of GABA_B_ antagonist (Serrano-Regal et al., 2020a). Whether the same is true during remyelination should be addressed in future using models with conditional ablation of GABA_B_Rs on OPCs and/or OLs.

Another potential mechanism underlying increased remyelination in Bac-treated mice could be the effect of Bac on microglia/macrophages, as these cells are important modulators of OPC differentiation (Miron et al., 2013). It has been shown that microglia-specific ablation of GABA_B1_Rs in mice impairs synaptic refinement by microglia, which leads to behavioral abnormalities (Favuzzi et al., 2021), thus suggesting that GABA_B_R modulation can modify microglial function. Our results show that Bac treatment did not alter microglia/macrophage (Iba1^+^ cells) numbers at 12 dpl. However, we do not know whether this treatment may have induced microglial/macrophage modifications that could favor remyelination without altering their recruitment or proliferation. While further research should address the exact effect of Bac on microglia/macrophages in the context of remyelination/demyelination, it appears clear that, at least in the LPC model, Bac administration improves remyelination, either directly by stimulating GABA_B_Rs in oligodendroglia, and/or by enhancing microglia/macrophage support of remyelination. In conclusion, our results suggest that Bac, a drug approved for spasticity treatment in MS patients, could be considered as a potential therapeutic candidate to improve remyelination. Because this drug is already administered to a subset of patients with MS, advanced imaging techniques for non-invasive measurement of myelin content/remyelination (Kolb et al., 2021) could be applied to evaluate whether patients treated with Bac indeed show evidence of enhanced remyelination. This could be an important first step in evaluating the suitability of this drug as a pro-remyelinating/neuroprotective agent in MS.

## Acknowledgements

We thank Dr. F. Kirchhoff for kindly providing us with PLP-DsRed mice. Dr. A. Palomino, Dr. T. Quintela-López and S. Marcos for their technical assistance. C. Luchena MSc for her advice in statistics, and Dr. L. Escobar for her expert assistance with Leica TCS STED SP8 laser scanning confocal microscope. Support provided by SGIker from the University of the Basque Country (UPV/EHU) (Animal Unit and Analytical and High-Resolution Microscopy in Biomedicine) is also gratefully acknowledged.

## SUPPLEMENTARY MATERIAL

**Supplementary Table 1.** GABAergic agonists and antagonists used in this study.

| Product | Reference | Supplier | Concentration<br>(*in vitro*) |
| --- | --- | --- | --- |
| GABA | A2129 | Sigma-Aldrich | 100 μM |
| Gabazine | SR-95531 | Sigma-Aldrich | 50 μM |
| Baclofen | 0796 | Tocris Bioscience | 100 μM |
| Muscimol | 0289 | Tocris Bioscience | 100 μM |

**Supplementary Table 2.** Antibodies used in this study for histology and immnufluorescence analysis.

| Antibody | Host | Dilution<br>(rat tissue) | Dilution<br>(mouse tissue) | Supplier | Reference |
| --- | --- | --- | --- | --- | --- |
| anti-GABAR<sub>B1</sub> | rabbit | 1:200 | 1:200 | Alomone labs | AGB-001 |
| anti-GABAR<sub>B2</sub> | rabbit | 1:200 | 1:200 | Alomone labs | AGB-002 |
| anti-APC<br>(clone CC1) | mouse | 1:200 | 1:200 | Calbiochem | #OP80 |
| anti-Olig2 | mouse | 1:200, 1:1000 | 1:500 | Millipore | #MABN50 |
| anti-MBP | chicken | - | 1:200 | Millipore | #AB9348 |
| anti-PDGFRα | rat | - | 1:300 | BD Biosciences | #558774 |
| anti-Iba1 | guinea pig | - | 1:200 | Synaptic systems | 234004 |
| anti-Nkx2.2 | mouse | - | 1:20 | Developmental Studies<br>Hybridoma Bank | #Q4818001B |
| anti-GFAP | rabbit | - | 1:100 | Millipore | #AB5804 |

**Supplementary Table 3.** Antibodies used in this study for western blot analysis.

| Antibody | Host | Dilution | Supplier | Reference |
| --- | --- | --- | --- | --- |
| anti-MAG | mouse | 1:500 | Santa Cruz | sc-376145 |
| anti-CNPase | mouse | 1:1000 | Sigma-Aldrich | #C5922 |
| anti-MBP | mouse | 1:1000 | Biolegend | #SMI 99 |
| anti-GAPDH | mouse | 1:1000 | Millipore | #MAB374 |
| anti-β-tubulin | mouse | 1:5000 | abcam | ab7291 |

**Supplementary Figure 1.**
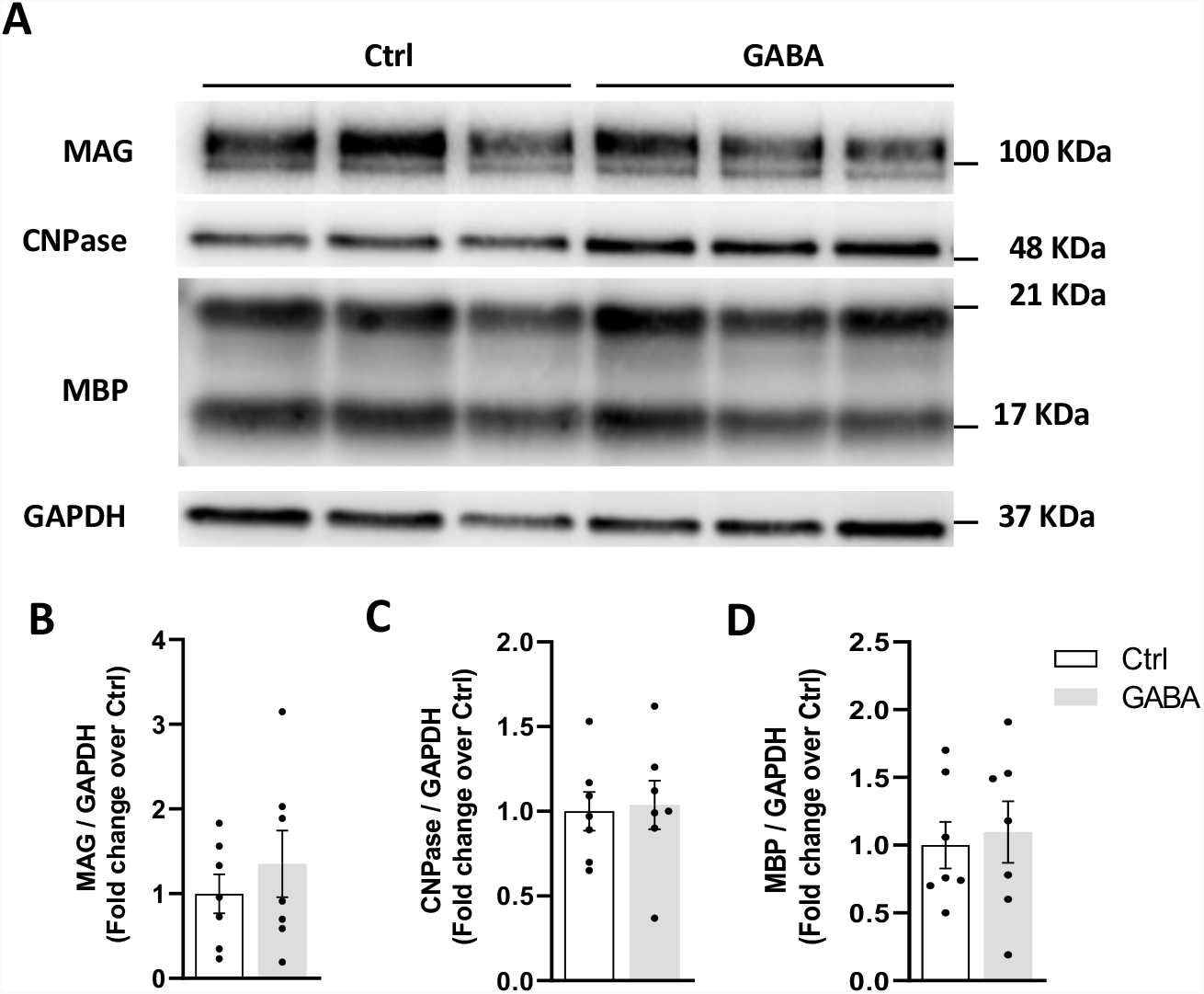
GABA addition does not change myelin-related protein levels in cerebellar organotypic slices. **A)** Western blot image showing expression of MAG, CNPase and MBP proteins in control condition or in presence of exogenous 100 μM GABA for 13 days in triplicates. Quantification of MAG **(B)**, CNPase **(C)** and MBP **(D)** expression normalized to GAPDH values. At least 3 independent experiments were included in the analysis.

**Supplementary Figure 2.**
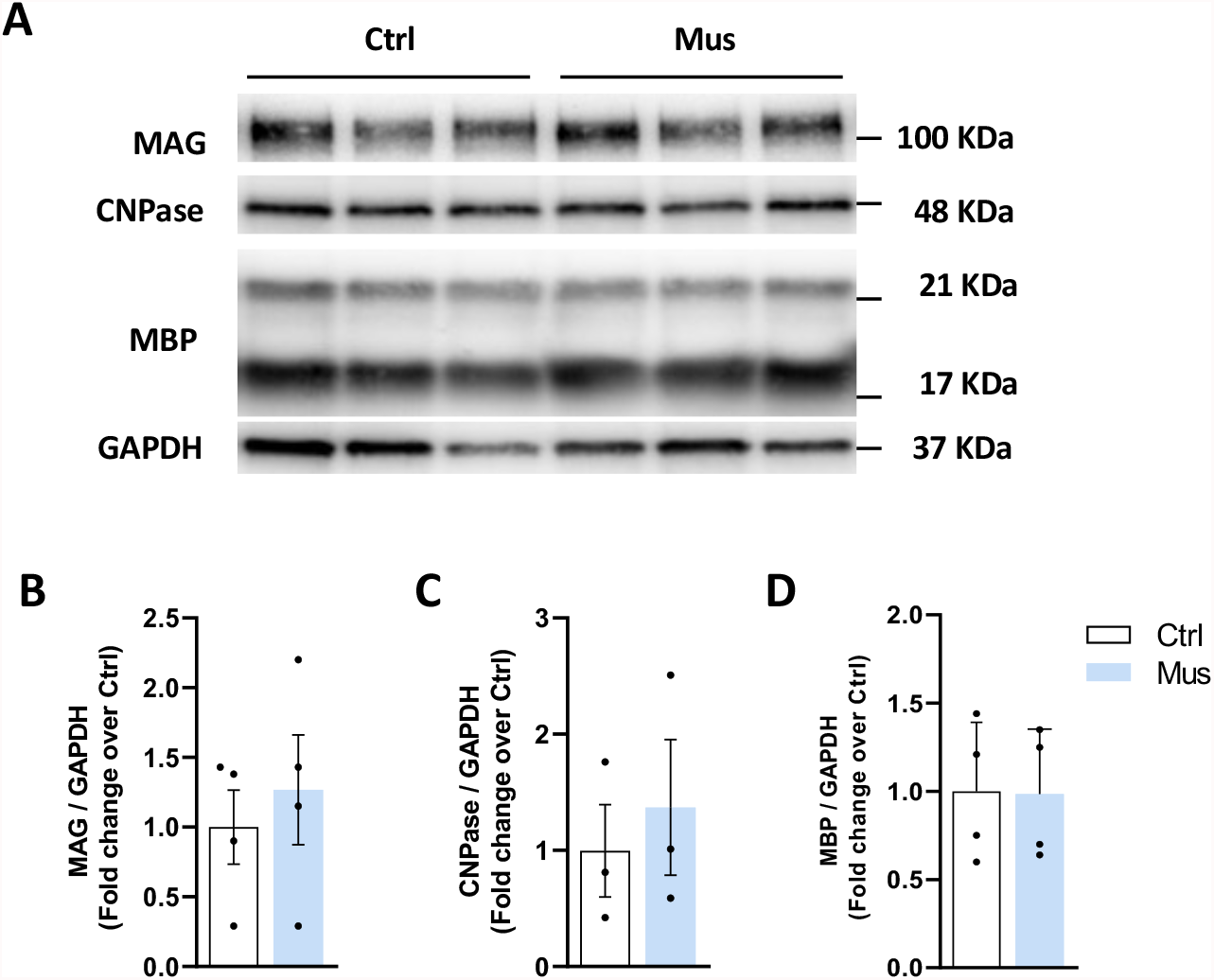
GABA_A_R stimulation does not modify the expression of myelin-related proteins in cerebellar organotypic slices. **A)** Representative western blot showing expression of MAG, CNPase and MBP proteins in control or muscimol-treated conditions for 13 days in triplicates. Histograms showing quantification of MAG **(B)**, CNPase **(C)** and MBP **(D)** levels. Values for each condition were obtained from at least 3 independent experiments and normalized to GAPDH.

**Supplementary Figure 3.**
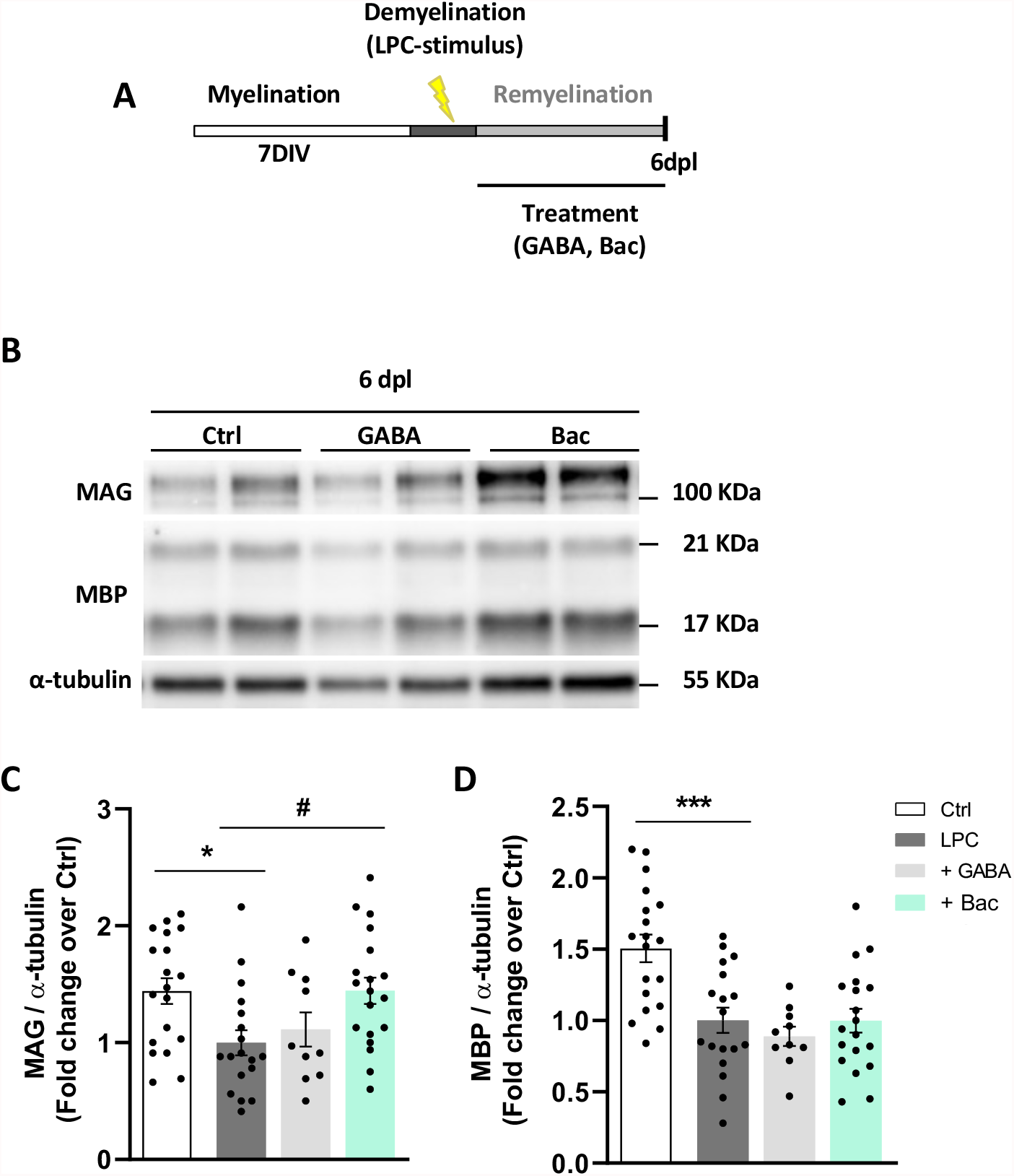
GABA_B_R activation increases MAG levels after LPC-induced demyelination in cerebellar organotypic cultures from P11 rats. **A)** Schematic representation of the experimental design of demyelination induced by LPC, treatment duration and the following remyelination. **B)** Representative western blot showing MAG and MBP levels after LPC exposure in duplicates. Values were obtained from at least 3 independent experiments and normalized to α-tubulin. Quantification of MAG **(C)** and MBP levels **(D)** in indicated conditions. *p<0.05 and ***p<0.001 *vs* control, ^#^p<0.05 *vs* LPC; one-way ANOVA followed by Tukey’s post-test.

**Supplementary Figure 4.**
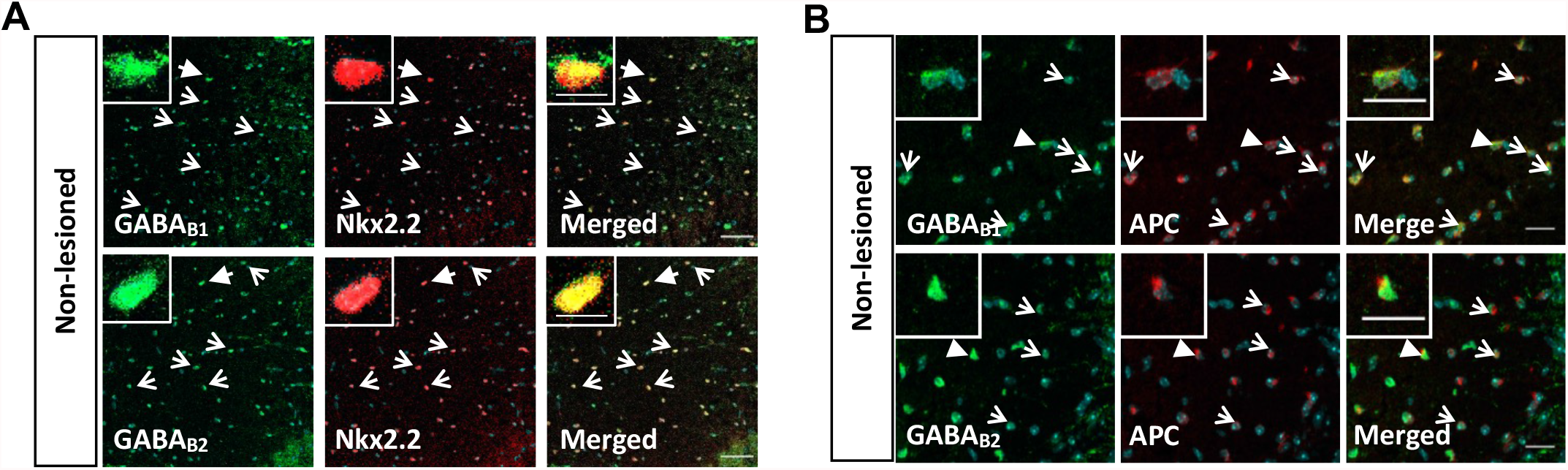
GABA_B_ receptors are present in OPCs and mature oligodendrocytes from the spinal cord of non-lesioned mice. **(A)** OPCs (Nkx2.2^+^, red) and **(B)** mature OLs (APC^+^, red) express GABA_B1_ and GABA_B2_ (green) subunits of GABA_B_Rs in the *dorsal funiculus* of the spinal cord of non-lesioned mice. Cells were identified by nuclear DAPI staining. Arrows indicate Nkx2.2^+^/APC^+^ cells expressing GABA_B_Rs subunits. Arrowheads point at cells shown at higher magnification in each photograph. Scale bars: **A** = 50 μm; **B** = 20 μm. Higher magnification = 10 μm.

